# Allosteric feedback inhibition of deoxy-D-xylulose-5-phosphate synthase involves monomerization of the active dimer

**DOI:** 10.1101/2022.06.12.495819

**Authors:** Xueni Di, David Ortega-Alarcon, Ramu Kakumanu, Edward E.K. Baidoo, Adrian Velazquez-Campoy, Manuel Rodríguez-Concepción, Jordi Perez-Gil

## Abstract

Isoprenoids are a very large and diverse family of metabolites required by all living organisms. All isoprenoids derive from the double-bond isomers isopentenyl diphosphate (IPP) and dimethylallyl diphosphate (DMAPP), which are produced by the methylerythritol 4-phosphate (MEP) pathway in bacteria and plant plastids. Understanding the regulation of the MEP pathway, probably the main metabolic pathway elucidated in this century, is a must for the rational design of biotechnological endeavors aimed at increasing isoprenoid contents in microbial and plant systems. It has been reported that IPP and DMAPP feedback regulate the activity of deoxyxylulose 5-phosphate (DXS), a dimeric enzyme catalyzing the main flux-controlling step of the MEP pathway. Here we provide experimental insights on the underlying mechanism. Our data show that direct allosteric binding of IPP and DMAPP to bacterial and plant DXS promotes monomerization of the enzyme. This allows a fast response to a sudden increase or decrease in IPP/DMAPP supply by rapidly shifting the dimer-monomer equilibrium accordingly. DXS monomers expose hydrophobic domains that are hidden in the dimer, resulting in aggregation and eventual degradation. Removal of monomers that would otherwise be available for dimerization and enzyme reactivation appears as a more drastic response in case of persistent IPP/DMAPP overabundance (e.g., by a blockage in their conversion to downstream isoprenoids). Our model provides a mechanistic explanation of how IPP and DMAPP supply can be adapted to changes in their demand and it also explains the changes in DXS protein levels observed after long-term interference of the MEP pathway flux.

**Significance Statement:** Isoprenoids are a vast family of organic compounds with essential roles in respiration, photosynthesis, photoprotection, membrane structure, and signaling. Many of them have great economic and nutritional relevance as pigments, aromas, drugs or phytonutrients. Despite their functional and structural diversity, they all derive from the same five-carbon precursors. We show that these precursors feedback-regulate their own synthesis in bacteria and plant plastids by allosterically shifting the dimer:monomer equilibrium of the enzyme that catalyzes the first step of their biosynthetic pathway towards the inactive monomeric form. This evolutionary conserved mechanism allows for both short-term (immediate) and long-term (sustained) control of the pathway flux, and its manipulation could be critical for the rational engineering of high-value isoprenoid products in bacterial and plant systems.

## Introduction

Isoprenoids (also known as terpenoids or terpenes) are a vast family of natural compounds produced in all free-living forms of life. Despite their astonishing variety both at the structural and functional levels, all isoprenoids derive from the universal building blocks isopentenyl diphosphate (IPP) and its isomer dimethylallyl diphosphate (DMAPP). In nature, two different pathways are responsible for the synthesis of IPP and DMAPP (1). The mevalonate (MVA) pathway is present mainly in eukaryotes and archaea and uses acetyl-CoA as an initial substrate whereas the methylerythritol 4-phosphate (MEP) pathway is found in most bacteria and produces IPP and DMAPP from pyruvate and glyceraldehyde 3-phosphate (GAP). In plants both pathways coexist but in different subcellular compartments. The MVA pathway produces IPP and DMAPP in the cytosol to synthesize sterols and sesquiterpenes, whereas the MEP pathway is located in plastids and produces the precursors of plastidial isoprenoids such as monoterpenes, carotenoids, and the side chains of chlorophylls, tocopherols, phylloquinones and plastoquinone.

The first step of the MEP pathway is catalyzed by the enzyme deoxyxylulose 5-phosphate (DXP) synthase (DXS), which uses pyruvate and GAP to generate DXP (Fig. 1). DXP is converted into MEP by the next enzyme of the pathway, DXP reductoisomerase (DXR). Metabolic control analyses showed that DXS is the enzyme with the highest flux control coefficient of the MEP pathway and consequently the primary rate-limiting step of the pathway both in bacteria and plants (2, 3). Consistent with this central role in the regulation of pathway flux, DXS activity is regulated at multiple levels including transcriptional, post-transcriptional and post-translational (1–8). Several reports have highlighted the importance of the metabolic regulation of the MEP pathway (1, 5, 9) Labeling experiments suggested a negative feedback regulation of plant DXS activity by MEP pathway products, IPP and DMAPP (10), which was later confirmed both *in vitro* and *in vivo* (2, 11–13). Interestingly, MEP pathway flux has been observed to also modulate DXS protein levels in plants (5, 12, 14). The feedback regulation of DXS levels and activity was shown to stabilize the plant pathway flux against changes in substrate supply and adjust it according to product demand under normal growth conditions (5). More recently, *in vitro* assays have shown a similar feedback regulation for several bacterial DXS enzymes (15), but *in vivo* confirmation is still missing.

**Fig 1.**
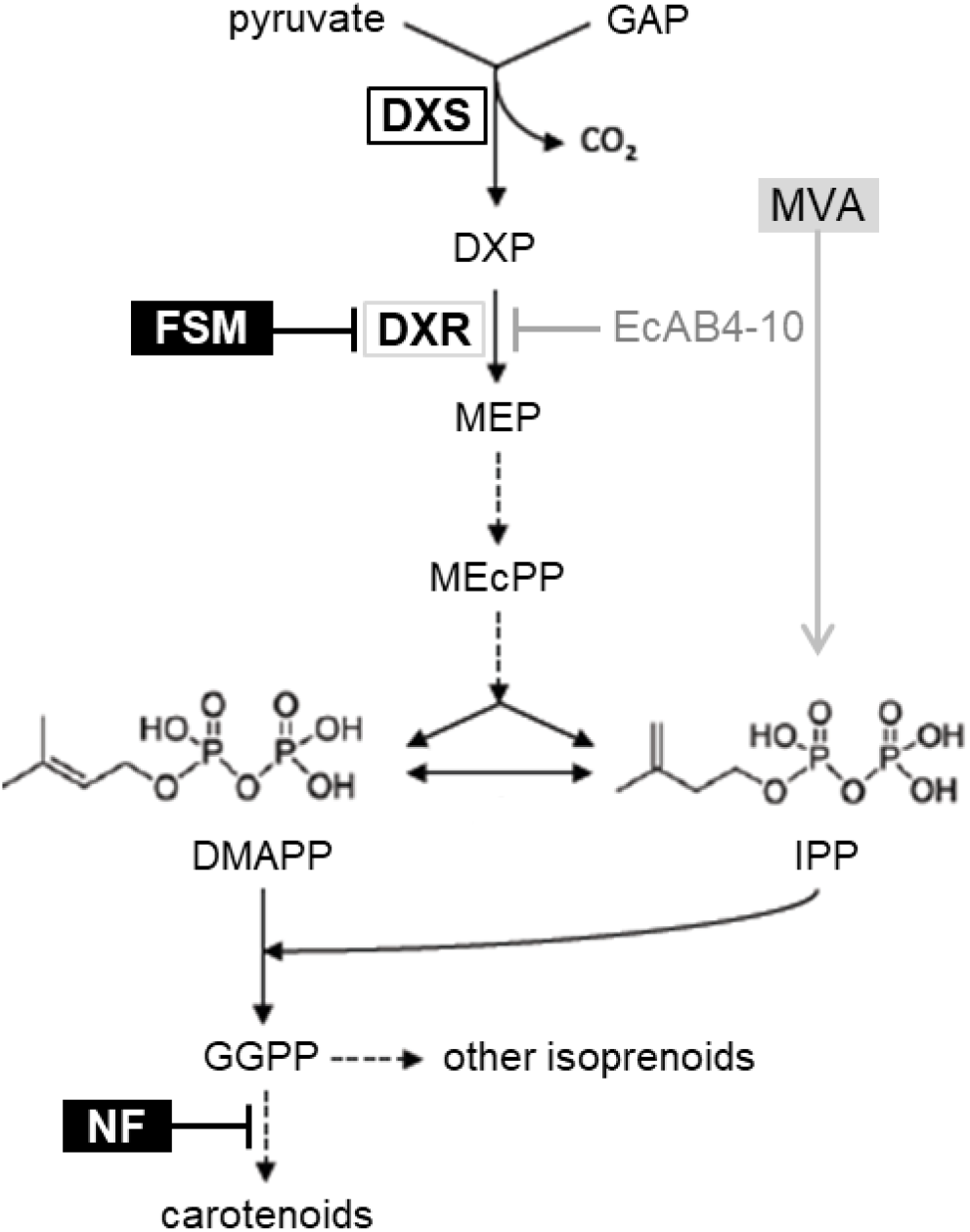
The MEP pathway and DXS. GAP, glyceraldehyde 3-phosphate; DXP, deoxyxylulose 5-phosphate; MEP, methylerythritol 4-phosphate; MEcPP, methylerythritol cyclodiphosphate; IPP, isopentenyl diphosphate; DMAPP, dimethylallyl diphosphate; GGPP, geranylgeranyl diphosphate; MVA, mevalonic acid. Enzymes are shown in bold: DXS, DXP synthase; DXR, DXP reductoisomerase. Inhibitors are boxed in black: FSM, fosmidomycin; NF, norflurazon. Dashed closed arrows represent several steps. The MEP pathway simultaneously produces both IPP and DMAPP for isoprenoid biosynthesis in bacteria and plant plastids. A synthetic operon transforming exogenously supplied MVA into IPP and DMAPP allows survival of MEP-defective *E. coli* strains such as EcAB4-10, which lacks DXR activity.

The initial crystallization of a truncated *Escherichia coli* DXS (16) was recently followed by reported structures of DXS enzymes from another bacterium, *Deinococcus radiodurans* (17), and from the model plant *Arabidopsis thaliana* (8). From these studies it was concluded that the active enzyme is a dimer and that a highly hydrophobic surface is present in the interface of the two monomers. Computational analysis of the monomer structure suggested that these hydrophobic domains increase the aggregation propensity of the protein (6). Arabidopsis DXS was actually found to easily aggregate, leading to subsequent degradation by the plastidial Clp protease complex. When stress conditions compromise the proteolytic capacity of the plastid, however, disaggregation rather than degradation is promoted for spontaneous refolding and reactivation of DXS (4, 6). The high aggregation propensity of DXS was also confirmed in cyanobacteria (18) and *E. coli* (15), whereas the involvement of the Clp protease in DXS degradation has also been described in tobacco *(Nicotiana tabacum)* leaves (19), tomato *(Solanum lycopersicum)* fruit (20), the malaria parasite *Plasmodium falciparum* (21) and *E. coli* (22).

Considering (a) that reduced IPP and DMAPP levels result in higher DXS activity and protein levels, (b) that DXS activity requires dimerization, (c) that monomers expose aggregation-prone hydrophobic domains, and (d) that aggregation normally leads to DXS degradation, we hypothesized that high IPP or DMAPP levels might displace the equilibrium towards the monomeric (i.e., inactive) conformation of the enzyme. If sustained, high IPP and DMAPP would lead to DXS monomer aggregation and eventual degradation. Here we tested this hypothesis using bacterial (*E. coli)* and plant (tomato) DXS enzymes.

## Results

### IPP/DMAPP inhibit EcDXS activity *in vivo*

Feedback regulation of DXS by MEP pathway products (IPP/DMAPP) has been described both *in vitro* and *in vivo* in plants (2, 10, 11) but only *in vitro* in bacteria (15). To validate the effect of IPP and DMAPP on the activity of *E. coli* DXS (EcDXS) *in vivo,* we used engineered strains harboring a synthetic MVA operon that allows the production of IPP and DMAPP when MVA is supplied to the growth medium (23) (Fig. 1). To accurately detect changes in DXS activity, we used the EcAB4-10 strain, which lacks DXR and hence it is unable to convert DXP into MEP and downstream IPP and DMAPP (23, 24). Survival of EcAB4-10 cells was allowed by supplementing the growth medium with MVA, which was converted into IPP and DMAPP by the synthetic MVA operon (23) (Fig. 1). EcAB4-10 cells were grown at 37°C in LB medium in the presence of 0.5 or 10 mM MVA and collected during the exponential phase to ensure steady-state conditions. Intracellular metabolites were then extracted and measured by LC-MS. IPP and DMAPP were quantified together due to the technical difficulty of separating these double-bond isomers. As expected, increasing amounts of intracellular MVA were measured in MVA-supplemented cultures, demonstrating the uptake of the molecule from the medium (Fig. 2). Also as expected, IPP/DMAPP levels were much increased in cells growing with 10 mM MVA compared to those growing with 0.5 mM MVA (Fig. 2). By contrast, DXP levels were much lower with 10 mM MVA, which is consistent with the conclusion that enhanced IPP/DMAPP levels cause a reduction of EcDXS activity in living cells (Fig. 2). It is important to note that EcAB4-10 cells growing with 10 mM MVA did not show any growth impairment compared to those growing with 0.5 mM MVA (Fig. 2), suggesting that the amounts of MVA-derived IPP/DMAPP were not reaching the very high levels previously reported to trigger toxicity (25, 26). Taken together, these results confirm that the negative feedback mechanism reported to control the activity of DXS also operates *in vivo* to decrease the activity of the bacterial EcDXS enzyme when IPP/DMAPP levels are increased.

**Fig 2.**
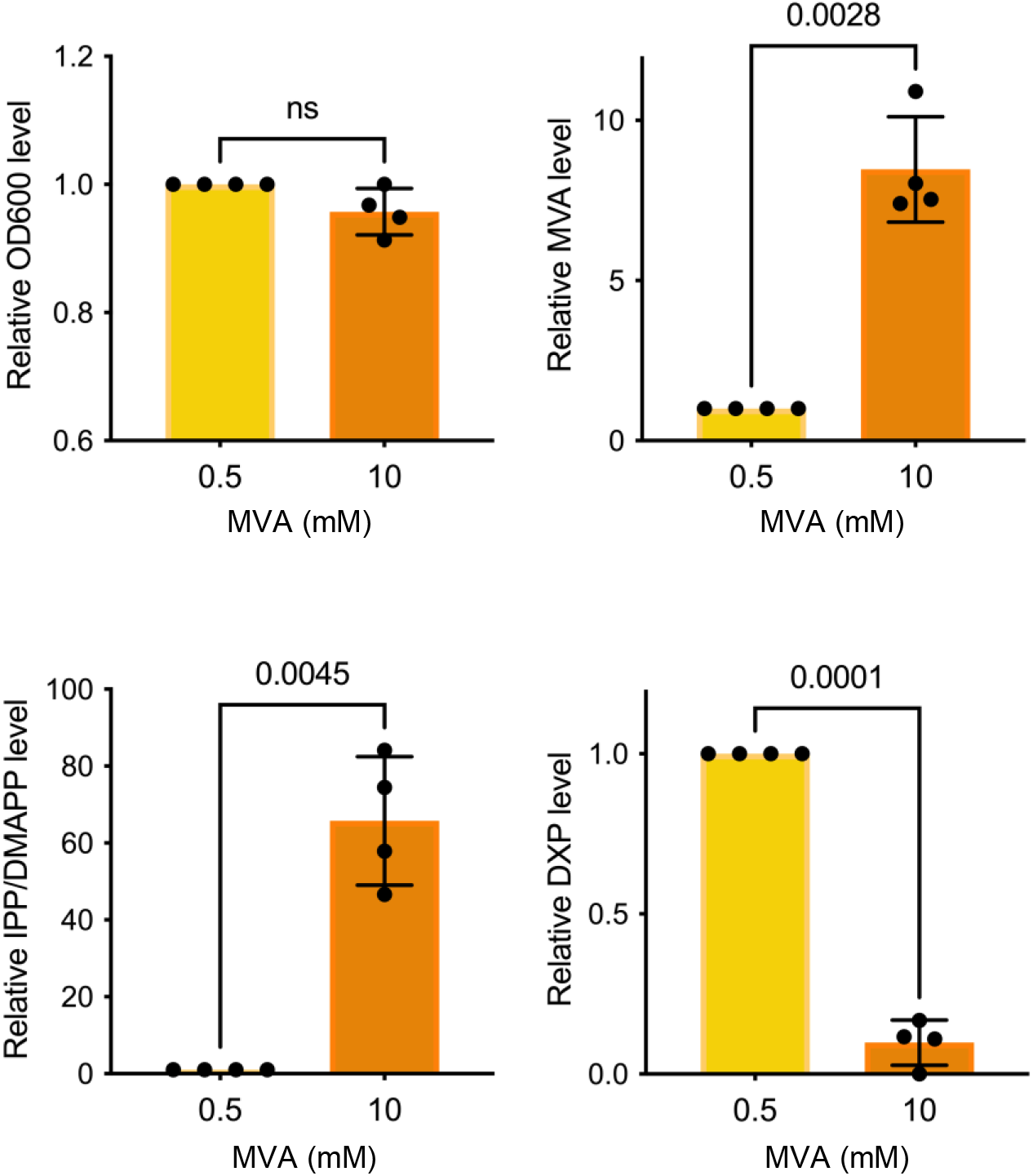
Metabolite levels in MVA-supplemented EcAB4-10 cells. *E. coli* cells of the EcAB4-10 strain were grown in the presence of 0.5 or 10 mM MVA and collected during exponential phase (steady-state) for metabolite extraction and quantification by LC-MS. Bar plots represent relative values of cell growth measured by its optical density at 600 nm (OD_600_) and intracellular levels of MVA, DXP and IPP/DMAPP. Dots represent individual values. Mean and standard deviation of the n=4 replicates are shown. The *p* values of Student t-test analyses are also indicated.

### IPP/DMAPP directly interact with DXS

Based on kinetic analyses and structural modeling, it was suggested that IPP and DMAPP might inhibit the activity of DXS by competing with its cofactor, thiamine diphosphate (TPP), for its binding site (11, 13). However, direct evidence is missing. In order to experimentally confirm this model, the interaction of DXS with IPP and DMAPP was assessed by isothermal titration calorimetry (ITC). Specifically, we tested purified 6xHis tagged versions of the full-length EcDXS protein or a truncated version of the tomato SlDXS1 isoform lacking the N-terminal plastid targeting peptide. However, the purified EcDXS protein was found to rapidly aggregate in solution and therefore we decided to only use the plant SlDXS1 enzyme for the ITC experiments together with TPP, IPP or/and DMAPP as ligands. Based on the thermograms obtained and the corresponding binding isotherms using 20 μM SlDXS1 in the sample cell solution and 200 μM of the ligand in the syringe solution, we confirmed the interaction with TPP, as expected, with a dissociation constant (K_d_) of 4.2 μM (Table 1 and Fig. S1). IPP and DMAPP were also found to interact with the plant enzyme, showing similar dissociation constants (K_d_ of 2.2 μM for IPP and 1.7 μM for DMAPP). Similar K_d_ values were also found when the SlDXS1 protein was preincubated with 100 μM TPP in the sample cell (K_d_ of 1.2 μM for IPP and 4.3 μM for DMAPP), suggesting the absence of competition for the TPP binding (i.e. active) site and consequently the existence of an unrelated binding site for IPP/DMAPP (Table 1 and Fig. S1). To explore whether binding of IPP and DMAPP had any effect on SlDXS1 structure, the protein was mixed with 50 μM TPP in the absence or presence of 50 μM IPP or DMAPP and then analyzed by dynamic light scattering (DLS). In the absence of IPP/DMAPP the estimated radius of the protein (5.8 nm) correlates with the dimeric form of the enzyme, of about 150 kDa. By contrast, in the presence of either IPP or DMAPP the peak shifted to a smaller radius while the range of particle sizes increased (Fig. 3). From these data we speculated that IPP and DMAPP binding might somehow interfere with the dimerization of SlDXS1, displacing the monomer-dimer equilibrium to the monomeric forms that might eventually aggregate forming the high molecular weight particles detected by DLS (Fig. 3).

**Table 1.** ITC-calculated association and dissociation constants.

| | | $K_a$ ($M^{-1}$) | $K_d$ ($\mu M$) |
| --- | --- | --- | --- |
| SIDXS1 | TPP | $2.3 \cdot 10^5$ | 4.2 |
| | IPP | $4.5 \cdot 10^5$ | 2.2 |
| | DMAPP | $5.8 \cdot 10^5$ | 1.7 |
| SIDXS1+TPP | IPP | $8.1 \cdot 10^5$ | 1.2 |
| | DMAPP | $2.3 \cdot 10^5$ | 4.3 |

**Fig 3.**
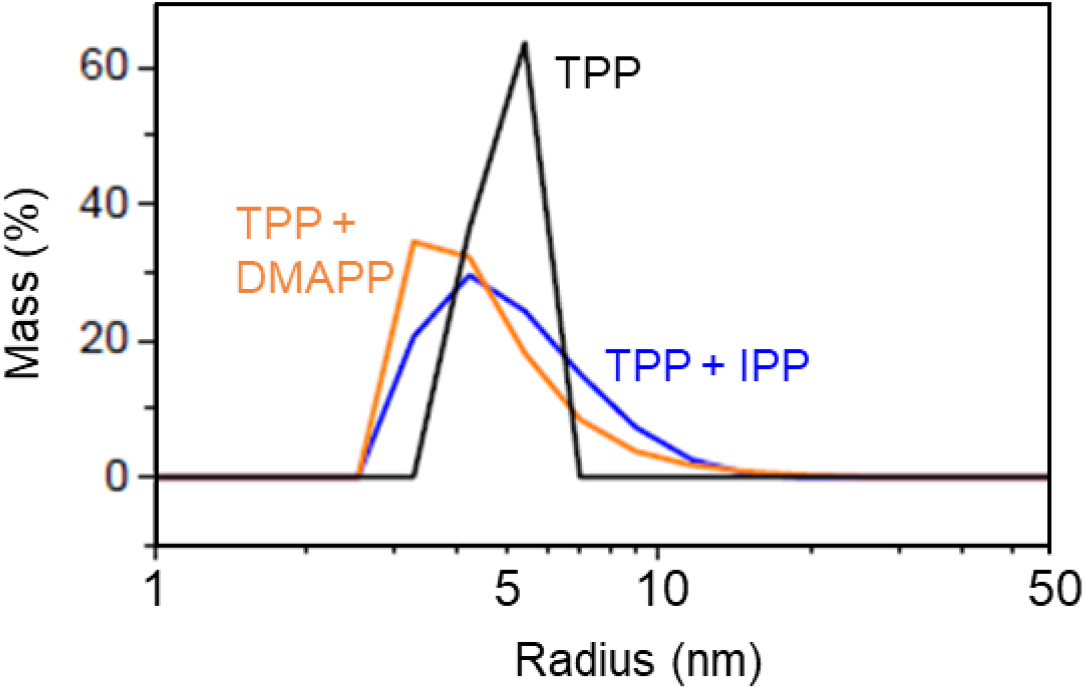
DLS analysis of purified SlDXS1 with ligands. Recombinant SlDXS (20 μM) was premixed with 50 μM TPP and analyzed in the absence (black line) or presence of 50 μM IPP (blue line) or 50 μM DMAPP (orange line). For each measurement, 5 acquisitions of 5 seconds were taken.

### IPP/DMAPP promote monomerization of bacterial and plant DXS enzymes

To further validate our interpretation of DLS experiments, we analyzed whether the oligomeric state of DXS enzymes from *E. coli* (EcDXS) and tomato (SlDXS1) changed in the presence of high levels of IPP/DMAPP *in vivo.* We initially expressed constructs encoding 6xHis-tagged versions of the full length EcDXS protein or the truncated SlDXS1 protein lacking the N-terminal plastid targeting peptide in the *E. coli* strain EcAM5, which harbors the MVA operon in a BL21(DE3) background (27). Transformed cells were grown at 37°C until exponential phase and then arabinose and IPTG were added to induce the expression of the MVA operon and the recombinant DXS enzyme, respectively. Cultures were also supplemented with 0, 1 or 10 mM MVA and then incubated for 3 additional hours at 26°C. The quaternary structure of recombinant DXS proteins in the cultures was analyzed by PAGE-SDS followed by immunoblot analysis with an anti-6xHis antibody after incubating total protein extracts with the crosslinker dimethyl suberimidate (DMS). In the presence of increasing concentrations of MVA, i.e., as IPP/DMAPP levels increased, the proportion of dimers decreased whereas monomeric forms of both EcDXS and SlDXS1 increased (Fig. 4). The structural change was even clearer when representing the dimer to monomer ratio (Fig. 4). These results demonstrate that high IPP and DMAPP levels decrease the proportion of active DXS dimers and increase the amounts of monomeric (inactive) forms of both bacterial and plant enzymes when expressed in *E. coli* cells.

**Fig 4.**
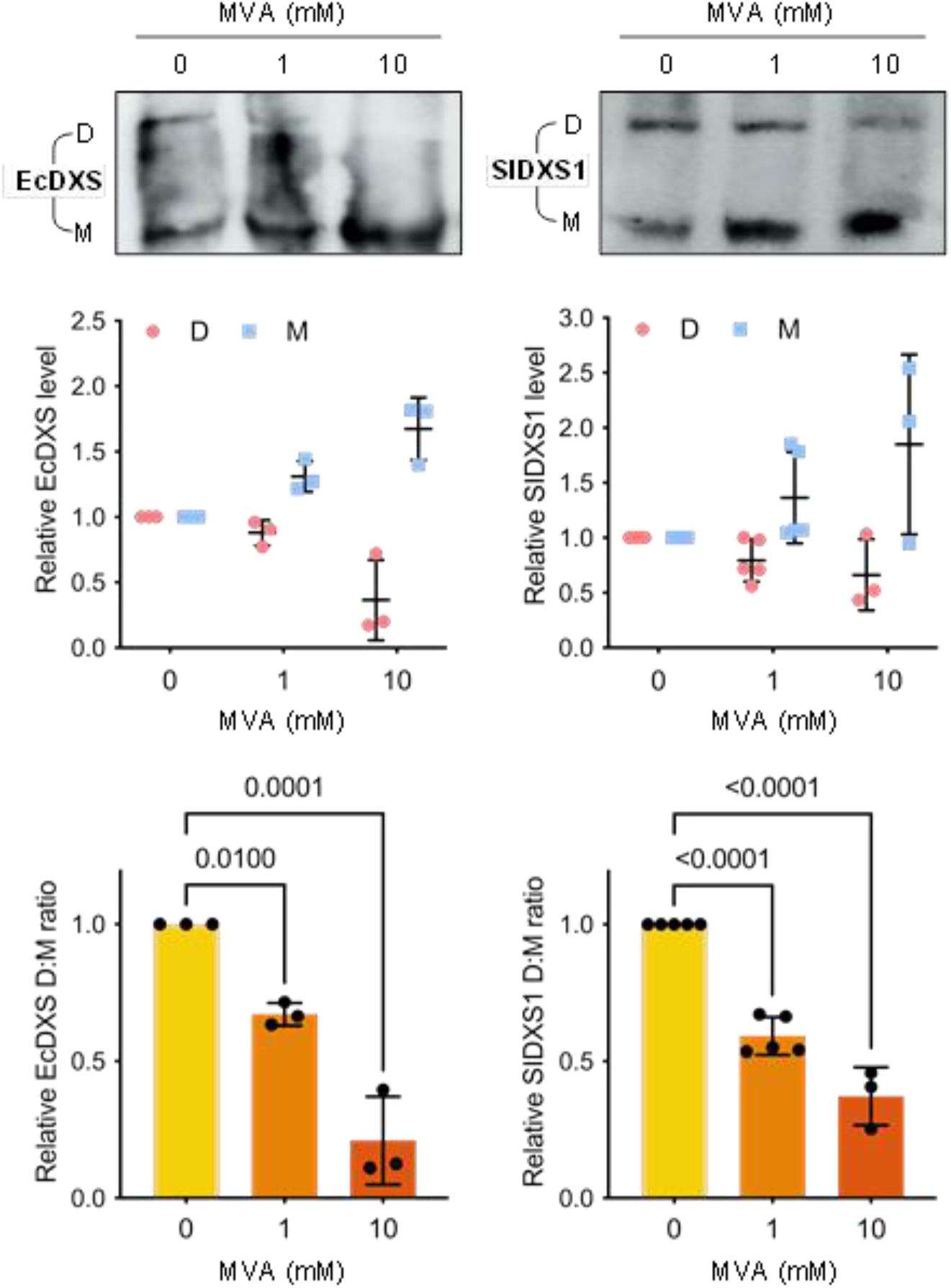
Increasing IPP/DMAPP levels promote monomerization of bacteria and plant DXS enzymes in *E. coli.* EcAM5<u>-1</u> cells were transformed with constructs to express 6xHis-tagged versions of bacterial (EcDXS) and plant (SlDXS1) enzymes and positive transformants were grown in media supplemented with the indicated amounts of MVA. Upper panels show representative images of immunoblot analyses with anti-6xHis antibodies. The position of dimers (D) and monomers (M) is indicated. Quantification of D and M abundance from immunoblot analyses of n≥3 independent experiments is represented in the plots. Individual data points as well as mean and standard deviation are represented. Numbers in the lower plots indicate *p* values (one-way ANOVA with Dunnett’s multiple comparisons test).

In order to validate the results obtained with the tomato SlDXS1 protein using a plant system, we agroinfiltrated constructs encoding the full-length protein in tobacco *(Nicotiana benthamiana)* leaves. The amount of IPP/DMAPP in leaf cells was modulated using the MEP pathway inhibitor fosmidomycin (FSM) and the carotenoid biosynthesis inhibitor norflurazon (NF) (Fig. 1). While both inhibitors cause a bleaching phenotype due to the photooxidative damage associated to the loss of carotenoids (Fig. 5A), FSM causes a drop in IPP/DMAPP levels whereas NF is expected to increase the amount of these isoprenoid precursors by preventing their consumption by the carotenoid pathway (Fig. 1). Leaf bleaching associated to the activity of these inhibitors, which were added to the agroinfiltration mixture, was clearly observed at 5 dpi (days post-inoculation) (Fig. 5A). This time point was selected to take samples for LC-MS analysis of MEP pathway intermediates as well as for protein extraction and immunoblot analysis using an anti-DXS serum (4). Our LC-MS method was not sensitive enough to detect IPP/DMAPP in agroinfiltrated tobacco leaves but we could successfully quantify the levels of DXP, MEP and the downstream intermediate methylerythritol-cyclodiphosphate (ME-cPP). FSM treatment caused an accumulation of DXP and a drop in MEP and MEcPP levels compared to untreated controls (Table 2). By contrast, NF treatment reduced the levels of DXP, MEP and MEcPP (Table 2), as expected considering that increased IPP/DMAPP levels should result in lower DXS activity. Consistent with our interpretation, the dimer to monomer ratio of SlDXS1 decreased in plant leaves treated with NF (Fig. 5B). Also in agreement with the results previously obtained in *E. coli*, reduced IPP/DMAPP levels in FSM-treated samples led to an increased dimer to monomer ratio compared to mock-treated controls (Fig. 5B). This negative correlation between dimer to monomer ratio and IPP/DMAPP levels strongly supports our conclusion that IPP and DMAPP are able to modulate DXS activity *in vivo* by interfering with the formation of the active dimeric form of the enzyme in both bacterial and plant systems.

**Fig 5.**
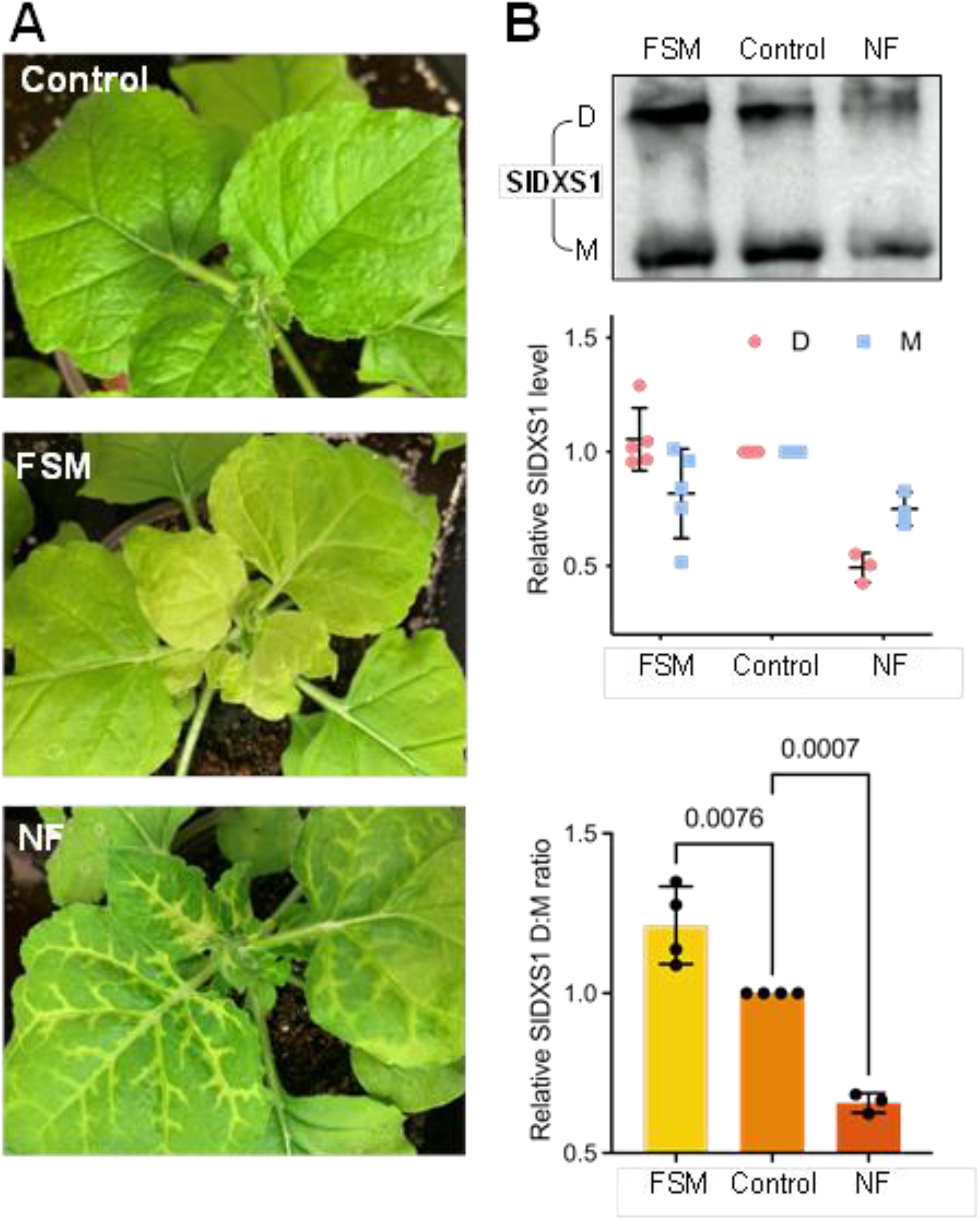
Changing IPP/DMAPP levels modulate monomerization of SlDXS1 in *N. benthamiana.* Leaves from *N. benthamiana* plants were infiltrated with constructs to produce SlDXS1 together with either water (control) or inhibitors, and samples were collected 5 days later. (A) Representative images of the leaves. (B) Immunoblot analyses of protein extracts with anti-DXS antibodies. The position of dimers (D) and monomers (M) is indicated in the upper panel, which shows the result of a representative experiment. Quantification of D and M abundance from immunoblot analyses of n≥3 independent experiments is represented in the plots. Individual data points as well as mean and standard deviation are represented. Numbers in the lower plots indicate *p* values (one-way ANOVA with Dunnett’s multiple comparisons test).

**Table 2.**
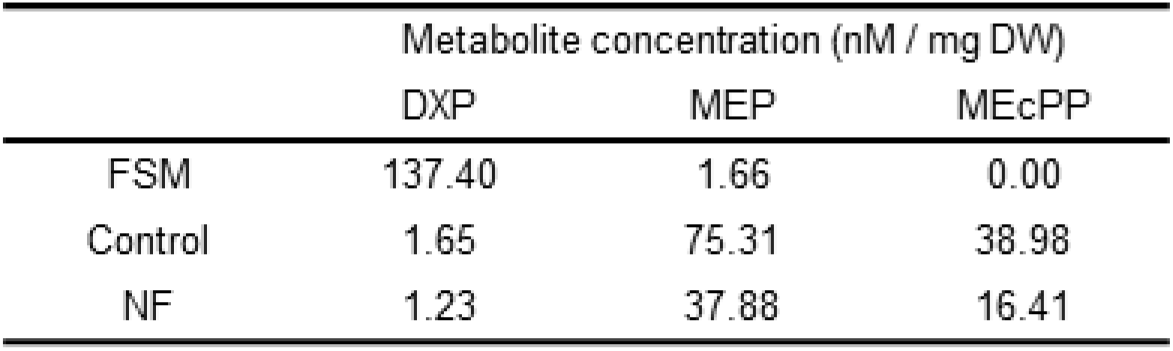
Concentration of MEP pathway intermediates in leaves infiltrated with the indicated inhibitors.

|  | Metabolite concentration (nM / mg DW) |  |  |
| --- | --- | --- | --- |
|  | DXP | MEP | MEcPP |
| FSM | 137.40 | 1.66 | 0.00 |
| Control | 1.65 | 75.31 | 38.98 |
| NF | 1.23 | 37.88 | 16.41 |

### IPP/DMAPP promote DXS aggregation in bacteria and plant cells

DXS monomers expose highly hydrophobic domains that make them prone to aggregation (6). To test whether IPP/DMAPP-induced monomerization of EcDXS and SlDXS1 proteins favored their aggregation, we expressed 6xHis-tagged versions of both proteins in EcAM5 cells, induced IPP/DMAPP accumulation *in vivo* by supplementing the growth medium with 1 or 10 mM MVA, and then isolated proteins from soluble and insoluble fractions for immunoblot analysis with anti-6xHis serum (Fig. 6). Most of the recombinant EcDXS and SlDXS1 proteins were found in the soluble fractions, with no major differences in the presence or absence of MVA (Fig. 6). In the insoluble fraction, however, the amounts of EcDXS and SlDXS1 increased in MVA-supplemented cells (i.e., when IPP/DMAPP levels increased), resulting in a higher ratio of insoluble (aggregated) vs. soluble (disaggregated) protein (Fig. 6).

**Fig 6.**
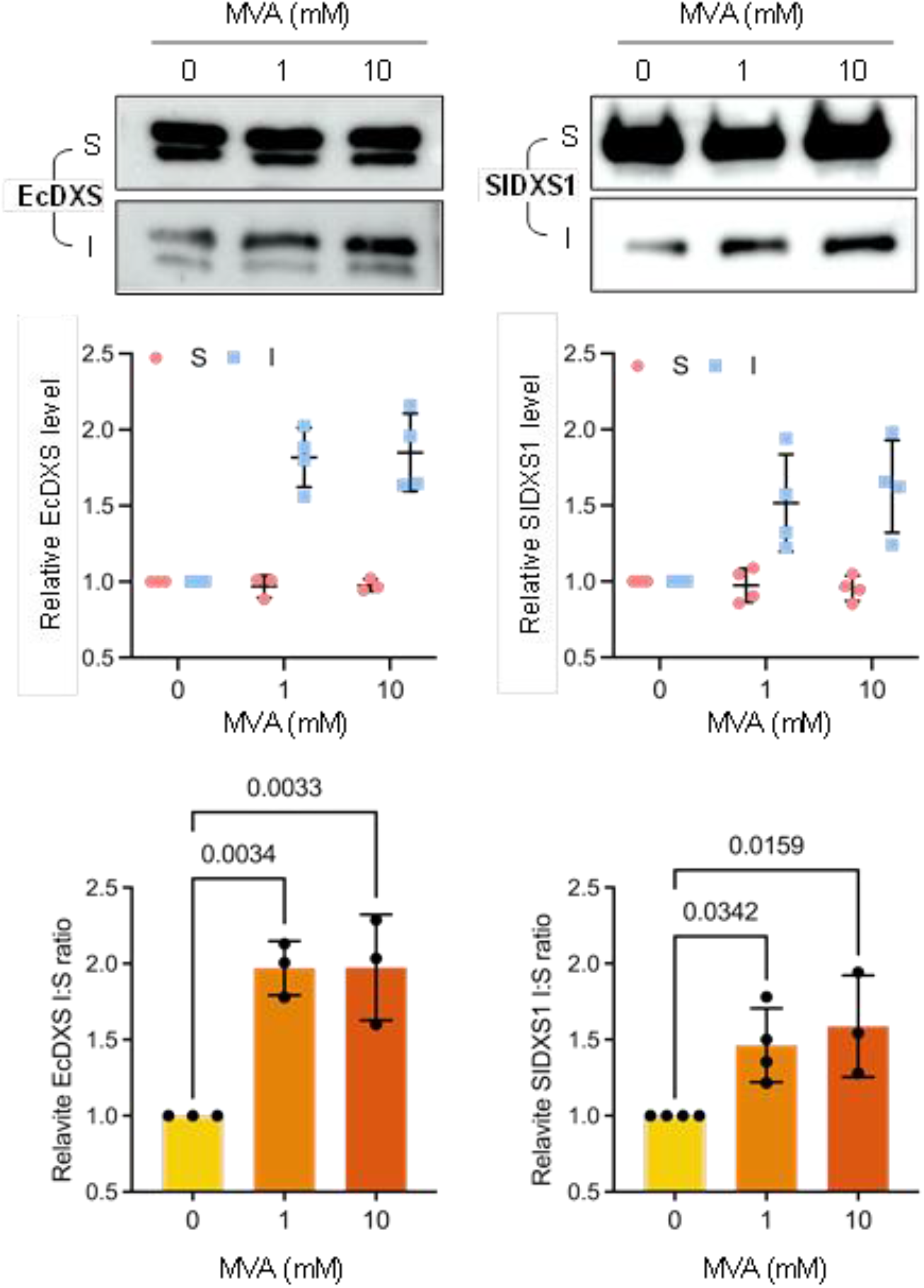
Increasing IPP/DMAPP levels promote aggregation of bacteria and plant DXS enzymes in *E. coli.* EcAM5 cells were transformed with constructs to express 6xHis-tagged versions of bacterial (EcDXS) and plant (SlDXS1) enzymes and positive transformants were grown in media supplemented with the indicated amounts of MVA. Upper panels show representative images of immunoblot analyses with anti-6xHis antibodies. Extracts corresponding to soluble (S) and insoluble/aggregated (I) proteins are indicated. Quantification of S and I abundance from immunoblot analyses of n≥3 independent experiments is represented in the plots. Individual data points as well as mean and standard deviation are represented. Numbers in the lower plots indicate *p* values (one-way ANOVA with Dunnett’s multiple comparisons test).

The aggregation status of the SlDXS1 protein was confirmed in plant cells by observing the accumulation of a GFP-tagged fusion, SlDXS1-GFP. Arabidopsis DXS enzymes fused to GFP are localized in the plastid stroma but they can also be observed forming a spotted distribution corresponding to aggregates when overexpressed (4, 6, 28). As expected, transient overexpression of SlDXS1-GFP in *N. benthamiana* leaves also led to the formation of fluorescent speckles identified by confocal laser scanning microscopy (Fig. 7). Quantification of the number of fluorescent spots (i.e. SlDXS1-GFP aggregates) per chloroplast in leaves treated with inhibitors altering IPP/DMAPP levels showed higher amounts with FSM and lower with NF compared to untreated controls (Fig. 7). The size of the spots, however, was higher in NF samples, in agreement with the conclusion that SlDXS1 aggregation increases when IPP/DMAPP levels increase.

**Fig 7.**
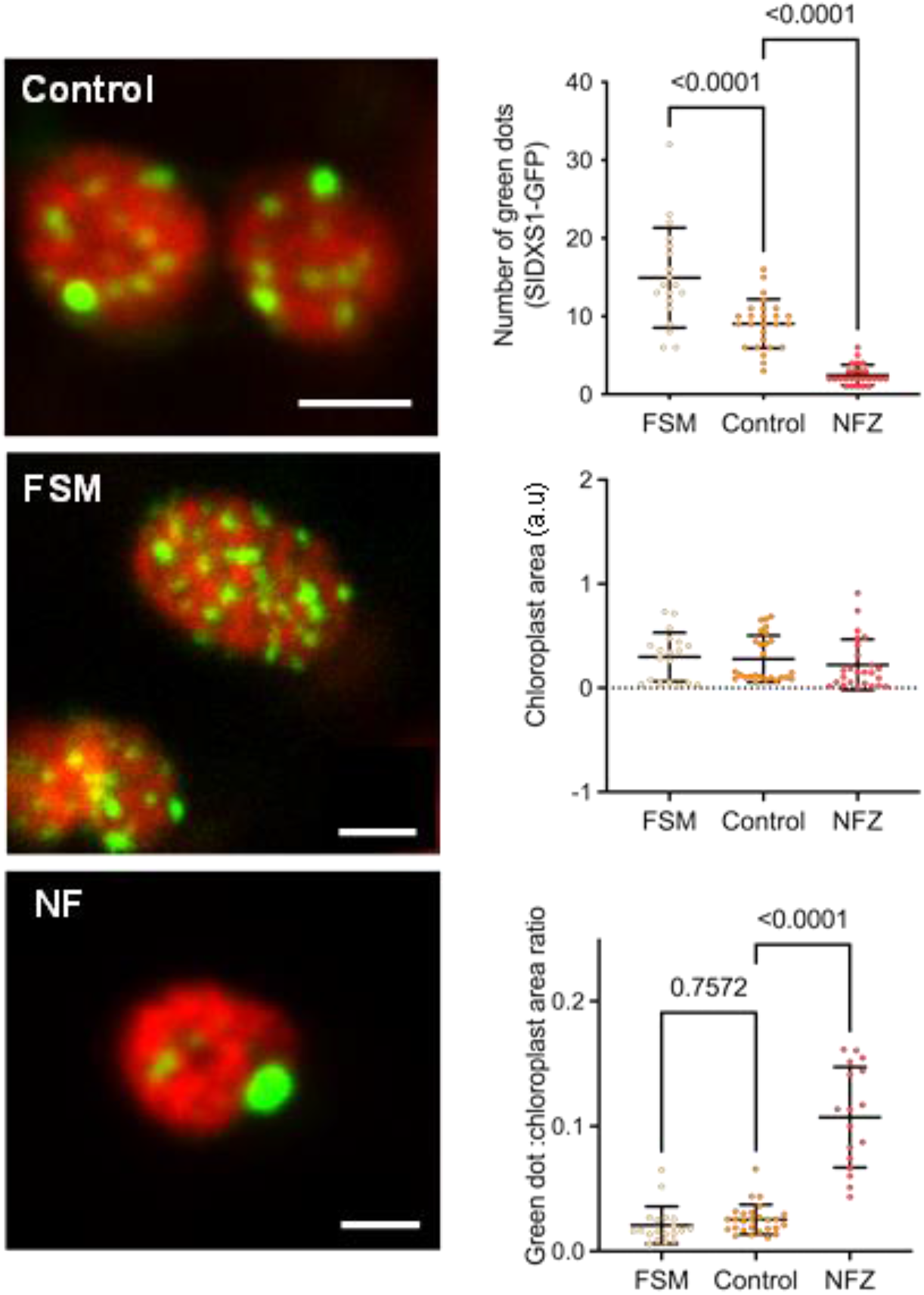
Changing IPP/DMAPP levels modulate aggregation of SlDXS1-GFP in *N. benthamiana.* Leaves from *N. benthamiana* plants were infiltrated with a constructs to produce SlDXS1-GFP together with either water (control) or inhibitors, and samples were analyzed 3 days later. Panels in the left show representative confocal microscopy images of SIDXS1-GFP (green) and chlorophyll (red) fluorescence in chloroplasts (size bar, 2 μm). Plots in the right show quantitative values from n≥7 pictures corresponding to different leaves. Individual data points as well as mean and standard deviation are represented. Numbers in the upper and lower plots indicate *p* values (one-way ANOVA with Dunnett’s multiple comparisons test). No significant differences in chloroplast area (middle plot) were found.

## Discussion

The regulation of the MEP pathway to produce the building blocks required for the biosynthesis of isoprenoids (IPP and DMAPP) has revealed to be rather complex, encompassing transcriptional, post-transcriptional, post-translational and metabolic levels (1, 9, 29, 30). This is especially relevant for the first enzyme of the pathway (DXS), which catalyzes the TPP-dependent conversion of pyruvate and GAP into DXP in the main rate-limiting step of the pathway (2, 3). Here we show that the mechanism by which accumulation of intracellular IPP and DMAPP inhibits the activity of DXS is conserved in bacteria and plant plastids and it involves (i) direct binding of IPP/DMAPP to DXS in a location different from the TPP-binding active site, followed by (ii) enhanced monomerization of the enzyme, and eventually (iii) aggregation of the inactive monomers (Fig. 8). This mechanism allows a short-term response to a sudden increase or decrease in IPP/DMAPP supply (by rapidly shifting the dimer-monomer equilibrium accordingly) but also a long-term response if IPP/DMAPP abundance persists (as monomers aggregate and become unavailable to form active DXS dimers).

**Fig 8.**
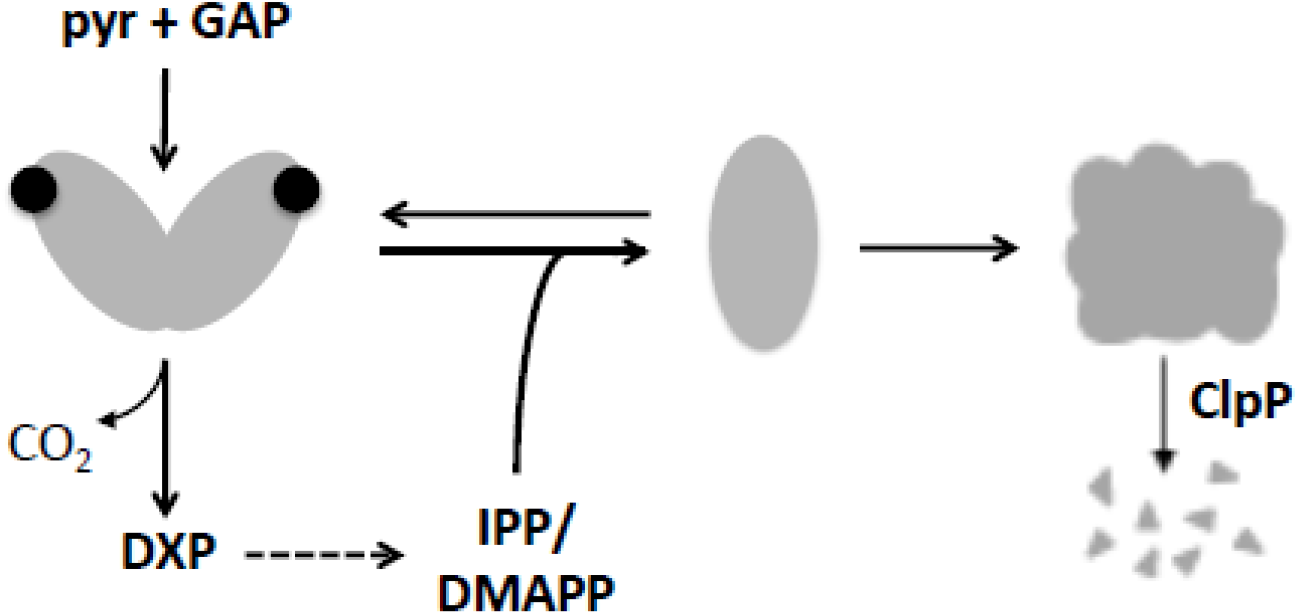
Model of the molecular mechanism involved in the allosteric feedback inhibition of DXS. The first step of the MEP pathway is the production of DXP from pyruvate and GAP, a reaction catalyzed by DXS. The active DXS enzyme is a dimer and requires binding of the cofactor TPP (represented as black circles) to the active site present in each of the subunits. Allosteric changes induced by binding of IPP/DMAPP shifts the dimer:monomer equilibrium towards the monomeric (inactive) forms, causing an immediate down-regulation of enzyme activity. If monomers are not used to make new dimers (e.g., due to the presence of high IPP/DMAPP levels), exposed hydrophobic domains can lead to monomer aggregation and eventual degradation by Clp protease complex (ClpP), resulting in a sustained removal of DXS activity.

DXS, in its dimeric active form, has been described as a three-domain polypeptide with domains I and II from the same chain involved in the formation of the active site and the binding of TPP. Domain III is involved in the formation of the dimer interface that includes a highly hydrophobic surface that remains unexposed to the solvent in the dimer (16, 17). The reaction mechanism involves the formation of a covalent intermediate between enzyme-bound TPP and pyruvate, followed by the GAP-stimulated decarboxylation of the bound pyruvate, and incorporation of GAP to the remaining fragment to generate DXP (16, 17, 31). Using a recombinant DXS protein from the tree *Populus trichocarpa* (PtDXS), it was proposed that IPP and DMAPP might compete with TPP for its pocket in the active site formed in each of the two subunits of the homodimer, with K_i_ values of ca. 65 μM for IPP and 81 μM for DMAPP (11). This competition is striking, however, considering that TPP is generally thought to be tightly bound as an integral part of DXS but also that it functions as a cofactor of many enzymes unrelated to isoprenoid synthesis for whom a regulation by IPP/DMAPP might not be physiologically meaningful. Experimental data from enzyme activity assays could not be fit to standard competitive inhibition kinetics and instead it was proposed a negative cooperative model in which binding of IPP/DMAPP to one of the DXS monomers would somehow hamper the binding of a second molecule to the other subunit of the homodimer (11). Furthermore, mutation of residues at the PtDXS active site that were predicted to be critical for binding the IPP/DMAPP carbon chain were found to have opposite effects on IPP-mediated inhibition of enzyme activity (13). Our ITC experiments using SlDXS1 in the presence of an excess of TPP, IPP and DMAPP confirmed that these three small diphosphate metabolites could indeed interact with the enzyme but showed K_d_ values for IPP and DMAPP that were very similar in the absence or presence of TPP (Table 1 and Fig. S1). These results argue against a direct competition of IPP/DMAPP for the TPP binding site in DXS and instead suggest the existence of an alternative binding site for the MEP pathway products. The inconsistencies between these two models of DXS inhibition by IPP/DMAPP might derive from the underlying methodology, i.e., enzymatic vs. calorimetric assays. ITC is a powerful tool to study protein/ligand binding interactions that involves directly measuring the heat that is released or absorbed in real time when one solution is titrated into another (32) ITC is capable of providing information on enzymatic reactions that is difficult to obtain using traditional (indirect) biochemical assays. In particular, the use of DXS substrates (pyruvate) together with additional metabolites (dihydroxyacetone phosphate) and enzymes (triose-phosphate isomerase from rabbit muscle) in the reaction mixtures previously used to determine the inhibition dynamics of IPP/DMAPP (11) might have resulted in interferences influencing the final interpretation of the data.

Allosteric regulation of protein function occurs when the binding of a molecule either activating or inhibiting the activity of the protein takes place away from the active site. This differs from competitive inhibition wherein the inhibitor binds to the active site and prevents the protein’s natural substrate gaining access. In contrast with the previously proposed model of negative cooperative inhibition of TPP binding by IPP/DMAPP (11, 13), our data supports the conclusion that IPP/DMAPP-mediated inhibition of DXS is truly allosteric as the modulation of enzyme activity involves binding outside the active site. Many proteins are allosterically regulated with a variety of mechanisms (33). Our model of IPP/DMAPP inhibition of DXS activity is a typical example of allosteric regulation in which the downstream products of a biosynthetic pathway down-regulate the activity of the enzyme catalyzing the first committed step through feedback inhibition, hence ensuring that pathway flux is adjusted to end-product usage (32, 34, 35). In most cases, binding of the ligand inhibits enzyme activity by promoting a conformational change (33). Because prediction of the energetics and mechanisms of protein conformational changes by ITC remains very challenging, we used DLS to initially address this possibility. Indeed, our results showed that IPP/DMAPP binding disrupts the dimeric structure of the active SlDXS1 enzyme, causing a peak shift towards smaller (monomeric) forms (Fig. 3). Further analysis of EcDXS and SlDXS1 dimeric and monomeric forms by immunoblot experiments confirmed a decreased dimer to monomer ratio when levels of IPP and DMAPP increased in living bacteria (Fig. 4) and plant cells (Fig. 5). Changes in the dimer:monomer ratio triggered by allosteric regulation has been observed in different proteins, including receptors (36), membrane translocators (37) and enzymes (38). In the case of DXS, however, IPP/DMAPP-mediated monomerization was found to be linked to increased aggregation both in bacteria (Fig. 6) and plastids (Fig. 7). DXS exists in a bimodal distribution of open and closed conformations. In the catalytic reaction mechanism described for DXS, the dimeric enzyme adopts a closed conformation when pyruvate binds the TPP-containing active site, and then changes to an open state upon GAP binding (17, 39, 40). In the open conformation, solvent-accessible areas are increased in different segments near the active site, exposing hydrophobic domains in the interface of the two monomers which are normally buried in the quaternary structure of closed dimer. This might partially explain the previously observed propensity to aggregation of this protein in plants (4, 6) and microbes (15, 18). Most interestingly, DXS monomers fully expose the aggregation-prone domains (6). We therefore conclude that high IPP/DMAPP levels shift the dimer:monomer equilibrium of DXS to monomeric forms for rapid down-regulation of enzyme activity. Monomers would be expectedly available for immediate dimerization and hence enzyme reactivation upon return of IPP/DMAPP to steady state levels. However, if IPP/DMAPP levels remain high (e.g., by a blockage in their consumption by downstream isoprenoid biosynthetic pathways), monomers would remain available for aggregation to drastically reduce DXS activity.

Our model, summarized in Fig. 8, not only provides a mechanistic explanation of how MEP-derived IPP and DMAPP supply can be adapted to changes in their demand but it can also explain the changes in DXS protein levels observed after long-term interference of the MEP pathway flux (5, 12, 14). In particular, genetic or pharmacological reduction of IPP/DMAPP levels cause an up-regulation of DXS protein levels without concomitant changes in gene expression (5, 12, 14), whereas mutants unable to produce carotenoids and hence expectedly accumulating higher IPP/DMAPP levels (similar to NF treatments) showed an opposite phenotype of reduced DXS protein levels (5). Under non-stressed conditions, DXS aggregates are degraded by the Clp protease complex in different organisms, including plants but also microorganisms (4, 6, 19–22). We therefore propose that a prolonged shortage of IPP/DMAPP might lead to mostly dimeric (i.e., soluble and enzymatically active) DXS whereas sustained IPP/DMAPP abundance would cause an enhanced monomerization and aggregation of the protein, eventually resulting in their efficient removal by the Clp protease (Fig. 8).

In summary, we discovered an evolutionary conserved and mechanistically simple system in which the MEP pathway products IPP and DMAPP (the universal precursors of all isoprenoids) allosterically control the activity of DXS (the main rate-limiting enzyme of the pathway) by modulating the equilibrium between dimeric (active) and monomeric/aggregated (inactive) forms of the enzyme. Peak demands of MEP pathway products (e.g., to produce monoterpenes in response to a pathogen attack or carotenoids upon a high light stress) would decrease the amount of IPP/DMAPP, promoting DXS dimerization and hence maximizing activity to increase pathway flux. By contrast, reduced IPP/DMAPP consumption causing the build-up of these metabolites would promote DXS monomerization, which could eventually result in aggregation and subsequent proteolytic removal of the protein (Fig 8). Overall, these results improve our surprisingly scarce knowledge of how the MEP pathway is regulated (5, 9, 41). The MEP pathway, probably the main metabolic pathway elucidated in this century (42), provides the precursors for a large diversity of isoprenoids with high added value from the industrial and nutritional points of view. Understanding its regulation, in particular that of its main enzyme, DXS, is therefore a must for the rational design of biotechnological endeavors aimed at increasing isoprenoid contents in microbial and plant systems.

## Materials and Methods

### Bacterial strains, plant material and growth conditions

*Escherichia coli* and *Agrobacterium tumefaciens* strains were grown in Luria broth (LB: 10 g/l bacto tryptone, 5 g/l yeast extract, and 10 g NaCl, plus 15 g/l bactoagar for plates) supplemented with antibiotics to a final concentration of 100 μg/ml ampicillin, 50 μg/ml kanamycin, 34 μg/ml chloramphenicol, 100 μg/ml rifampicin, 30 μg/ml gentamicin, or/and 100 μg/ml spectinomycin, when required. Bacterial growth in liquid media was monitored by measuring optical density at 600 nm (OD_600_). The strains used in this study are described in Table S1. *N. benthamiana* plants were grown under standard greenhouse conditions.

### Gene constructs

To obtain 6xHis-tagged proteins, we cloned DXS-encoding bacterial and plant sequences into the pET23 vector (Novagen). The sequence of the *E. coli* gene encoding EcDXS was PCR-amplified from the wild-type strain MG1655 genomic DNA with primers EcDXS-NdeI-F and EcDXS-XhoI-R (Table S2) and the fragment obtained was digested with N*deI* and X*hoI* for ligation to a pET23 backbone (previously digested with the same enzymes) to generate the construction pET23-EcDXS. The coding sequence of SlDXS1 was amplified from a *S. lycopersicum* ripe fruit cDNA library using primers NheI-myc-SlDXS1-F and SlDXS1-XhoI-R (Table S2). The DNA fragment of the expected size was digested with *Nhe*I and *Xho*I and ligated to a pET23 vector previously digested with the same enzymes to yield the construct pET23-SlDXS1 for expression in *E. coli*. Constructs pGWB420-SlDXS1 and pGWB405-SlDXS1-GFP for expression in plants were obtained following a two-step (BP/LR) Gateway reaction yielding proteins fused to C-terminal myc or GFP tags, respectively. Primers and constructs are listed in Table S2. All constructs were sequenced to confirm the identity of the genes and the absence of undesired mutations.

### Production and purification of recombinant proteins

Following the transformation of *E. coli* BL21(DE3) pLysS cells with the required construct, the production of recombinant DXS proteins were induced by adding 50 μM IPTG to cultures with an OD_600_≈0.6. After growth for 14 h at 28 °C, bacterial cells were recovered by centrifugation and the recombinant protein was purified. The cell pellet was resuspended in Buffer A (50mM HEPES, 150 mM NaCl, pH=7.5) supplemented with 1 mg/mL lysozyme, 0.5 mM EDTA, and one tablet of complete protease inhibitor cocktail (Roche) for every 10 mL of buffer. The resuspended pellet was incubated on ice for 20 min with light shaking and after a brief sonication (6 pulses of 30 sec at 10% with 45 sec pause between pulses) the cell lysate was centrifuged at 27,000 *xg* for 20 min to separate broken cells. A 1:7 volume of protamine sulphate (1% in water) was added to the supernatant and the mixture was centrifuged at 39,000 x*g* for 1h. The cleaned and filtered supernatant was loaded into a poly prep chromatography column (BioRad) containing 1,5 ml (CV) of Ni-NTA agarose (Qiagen) previously cleaned and equilibrated with buffer A. After adding 10 mL of buffer A and 5 mL of washing buffer (buffer A containing 10 mM imidazole), the recombinant protein was eluted with 1.5 mL aliquots of elution buffer (buffer A supplemented with150 mM imidazole). Fractions containing the purified protein were pooled and imidazole removed using a PD10 desalting column (Cytiva) equilibrated with buffer A. The protein was collected after centrifugation at 1.000 x*g* for 2 min and stored as single-use aliquots at −80°C.

### *In vivo* inhibition assays

*E. coli* strain EcAB4-10 was grown on LB plates supplemented with kanamycin, chloramphenicol and MVA. Five colonies were grown in liquid medium at 37°C and 200 rpm o/n to inoculate a 15 ml fresh culture containing 0.1% arabinose and either 0.5 or 10 mM MVA at an initial OD_600_=0.1. Cells were grown at 37°C and 200 rpm until OD was ≈1 to ensure steady state for proper analysis, collected by centrifugation and stored at −80°C. For metabolite analysis, samples were thawed on ice, immediately quenched with 400 μl methanol and lysed mixing thoroughly. Then, 400 μl of water were added and mixed before centrifugation of the mixture at 13.000 x*g* for 5 min at 4°C. About 750 μl of the supernatant were collected and macromolecules were removed using a 3 kDa Amicon filter at 4°C and 13.000 x*g* for 1 hour. Then, 700 μl of the filtrate were mixed with 1 ml of water and flash-freezed in liquid nitrogen. Frozen samples were lyophilized and stored at −80°C until reconstitution with 100 μl of methanol:water (1:1) for LC-MS/MS analysis as described (44).

### Isothermal titration calorimetry (ITC)

The interaction between the DXS proteins and TPP, IPP or DMAPP was assessed using a high precision Auto-iTC200 calorimeter (MicroCal, Malvern-Panalytical). Protein solutions in the calorimetric cell at 20 μM were titrated with the different ligands (TPP, IPP, or DMAP) at 200 μM in the injecting syringe using buffer A (50 mM HEPES, 150 mM NaCl, pH=7.5) as the common buffer solvent. A sequence of 19 injections of 2 μL each was programmed, with a stirring speed of 750 rpm and 150 s spacing, and applying a reference power of 10 μcal/s. The association constant (K_a_) was estimated through non-linear least squares regression analysis of the data by using a model considering a single ligand binding site (1:1 protein:TPP/IPP/DMAPP) employing a user-defined fitting routine implemented in Origin 7.0 (OriginLab Northampton, MA). The dissociation constant (K_d_) was calculated as the inverse of K_a_.

### Dynamic light scattering (DLS)

Dynamic light scattering measurements were performed in a DynaPro Plate Reader III (Wyatt Technology) using a 384-multiwell plate (Aurora Microplates). Hydrodynamic radius of SlDXS1 was measured before and after the addition of TPP, DMAPP and IPP to estimate the size distribution of the protein. A second measurement was performed after 10 min incubation in order to assess the evolution of the complex. For each measurement, 5 acquisitions of 5 seconds were taken, and the apparent hydrodynamic radius was estimated from the experimental diffusion coefficient, obtained by the cumulant fit of the translational autocorrelation function, assuming an equivalent Rayleigh sphere model. Experiments were performed with a fixed protein concentration of 20 μM, and ligands IPP, DMAPP or TPP at 50 μM in the same buffer used for ITC experiments.

### *In vivo* DXS oligomeric state analysis in bacteria

*E. coli* EcAM5-1 cells were transformed with constructs pET23-EcDXS or pET23-SlDXS1 and single colonies were grown o/n in LB medium supplemented with ampicillin and kanamycin at 37°C. Fresh medium (150 ml) was inoculated at 1.5% and grown at 37°C and 200 rpm until OD_600_ ≈ 0.6. Following the addition of 0.1% arabinose and 0.05 mM IPTG, the culture was split in three and 0 mM, 1 mM or 10 mM MVA was added to each of the three subcultures. All subcultures were then grown at 26°C and 200 rpm for 3h and cells were collected by centrifugation. Samples were ground in liquid nitrogen and extracted with 0.2 M triethanolamine (pH=8). The mixture was centrifuged at 13.000 *xg* for 10 min at 4°C and the supernatant was treated for 40-50 min at room temperature with 10-fold molar excess of dimethyl suberimidate (DMS). The crosslinking reaction was stopped by adding 20 mM Tris. To separate soluble and insoluble (aggregate) protein fractions, samples were ground in liquid nitrogen, extracted with 0.2 M triethanolamine (pH=8), centrifuged at 13.000 x*g* for 10 min at 4°C, and the supernatant collected as the soluble protein fraction. The cell pellet was washed with 0.2 M triethanolamine and resuspended by insoluble protein extraction buffer (8M urea, 10mM Tris and 100mM NaH2PO4) for 20 min at room temperature. After centrifuging at 13.000 x*g* for 10 min, the supernatant was collected as the insoluble protein fraction. The concentrations of both soluble and insoluble protein fractions were determined using the Bio-Rad protein assay. Samples were resolved in a 7% SDS-PAGE, transferred to a PVDF membrane (Amersham), and used for immunoblot analysis with 1:2500 anti-6xHis antibody (Proteintech) for 10 min at room temperature. Detection of immunoreactive bands was performed using SuperSignal West Pico PLUS (Thermo Scientific). Chemiluminescent signals were visualized using an ImageQuan 800 biomolecular imager (Amersham) and quantified using Image J software. Statistical significance of quantified differences was analyzed with GraphPad.

### *In vivo* DXS oligomeric state analysis in planta

*Agrobacterium tumefaciens* GV3101 cells were transformed with the constructs pGWB420-SlDXS1 or pGWB405-SlDXS1-GFP and a single transformed colony was grown o/n in 3 ml LB medium supplemented with rifampicin, gentamycin and spectinomycin at 28°C. Fresh medium (25 mL) was inoculated with the preculture and grown o/n at 28°C. The culture was centrifuged to collect cells and the pellet was resuspended in agroinfiltration buffer (10 mM MgCl_2_, 10 mM MES, 150 μl acetosyringone, pH=5.6) to achieve OD_600_=0.8. The mixture was incubated at 28°C and 200 rpm for 1.5h and then mixed with a similar culture harboring vector pGWB702-HCProWMV to prevent silencing (43). The 9:1 (DXS:HcProWMV) mix was split in three and the individual fractions were supplemented with either fosmidomycin (200 μM), norflurazon (40 μM) or water before infiltration of leaves from 4-week-old *N. benthamiana* plants as described (43). Leaf samples agroinfiltrated with the pGWB420-SlDXS1 construct were collected at 5 dpi, snap-frozen in liquid nitrogen, and stored at −80°C. To extract proteins, around 300 mg of frozen tissue was ground in liquid nitrogen and mixed with 450 μl TKMES buffer (4) supplemented with 20 μl/ml protease inhibitor cocktail and 20 μl/ml 10% Triton-100 (v/v). The mixture was centrifuged at 2,800 x*g* for 10 min at 4°C and the supernatant was further cleared with two additional centrifugation steps. Protein concentration of the extract was measured with the Bio-Rad protein assay. Samples were mixed with non-denaturing loading buffer, separated in a 7% SDS-PAGE and then transferred to a PVDF membrane. The membrane was incubated overnight at 4 °C with 1:500 anti-DXS antibody (4). Chemiluminescent signals and statistical analysis were performed as described above. Subcellular localization of GFP-tagged SlDXS1 was observed by direct examination of agroinfiltrated leaf tissue at 3 dpi using confocal microscope AxioObserver 780 (Zeiss) with a BP515-525 filter after excitation at 488 nm. The areas of the whole chloroplast and the SlDXS1-GFP green fluorescent dots was quantified from pictures using Image J.

## Acknowledgments

This work was funded by grants from Spanish MCIN/AEI/10.13039/501100011033 and European ERDF/FEDER, NextGeneration EU/PRTR and PRIMA programs to MR-C (PID2020-115810GB-I00 and UToPIQ-PCI2021-121941) and AV-C (BFU2016-78232-P). MR-C is also supported by CSIC (202040E299) and Generalitat Valenciana (PROMETEU/2021/056). RK and EEKB conducted metabolite analysis at the Joint BioEnergy Institute (http://www.jbei.org), supported by the US Department of Energy, Office of Science, Office of Biological and Environmental Research under contract no. DE-AC02-05CH11231 between Lawrence Berkeley National Laboratory and the US Department of Energy. J.P-G was supported by a Marie Curie International Outgoing Fellowship within the EC-FP7 Program (Project Number 627639). X.D was supported by a China Scholarship Council and D.O-A by a MCIN/AEI/ fellowship (BES-2017-080739).

## Supplemental Tables

**Supplemental Table S1:** Strains used in this study.

| Name | Description | Antibiotic | Reference |
| --- | --- | --- | --- |
| TOP10 | F- *mcrA* Δ(*mrr-hsdRMS-mcrBC*) Φ80*lacZ*Δ*M15* Δ *lacX74* *recA*<sup>-</sup><br>*araD139* Δ(*araleu*)7697 *galU galK rpsL* (Str<sup>R</sup>) *endA1 nupG* | - | Invitrogen |
| BL21(DE3)pLysS | F<sup>-</sup> *ompT hsdSB* (r<sub>B</sub><sup>-</sup>, m<sub>B</sub><sup>-</sup>) *gal dcm rne131* (DE3) pLysS (Cm<sup>R</sup>) | Cm | Invitrogen |
| K12 MG1655 | F- *lambda- ilvG - rfb -50 rph -1* | - | (1) |
| EcAB4-10 | K12 MG1655 Δ*dxr::CAT* pBAD-yPMD-hPMK-yMVK-EcIDI | Km/Cm | (2) |
| EcAM5-1 | BL21(DE3) pBAD-yPMD-hPMK-yMVK-EcIDI | Km | (3) |

**Cm:** Chloramphenicol, **Km:** Kanamycin
1. F. R. Blattner *et al.*, The complete genome sequence of *Escherichia coli* K-12. *Science (New York, N.Y.)* **277**, 1453-1462 (1997). 2. S. Sauret-Güeto, E. M. Urós, E. Ibáñez, A. Boronat, M. Rodríguez-Concepción, A mutant pyruvate dehydrogenase E1 subunit allows survival of *Escherichia coli* strains defective in 1-deoxy-D-xylulose 5-phosphate synthase. *FEBS letters* **580**, 736-740 (2006). 3. A. Rodríguez-Villalon, J. Perez-Gil, M. Rodríguez-Concepción, Carotenoid accumulation in bacteria with enhanced supply of isoprenoid precursors by upregulation of exogenous or endogenous pathways. *Journal of biotechnology* **135**, 78-84 (2008).

**Supplemental Table S2:**
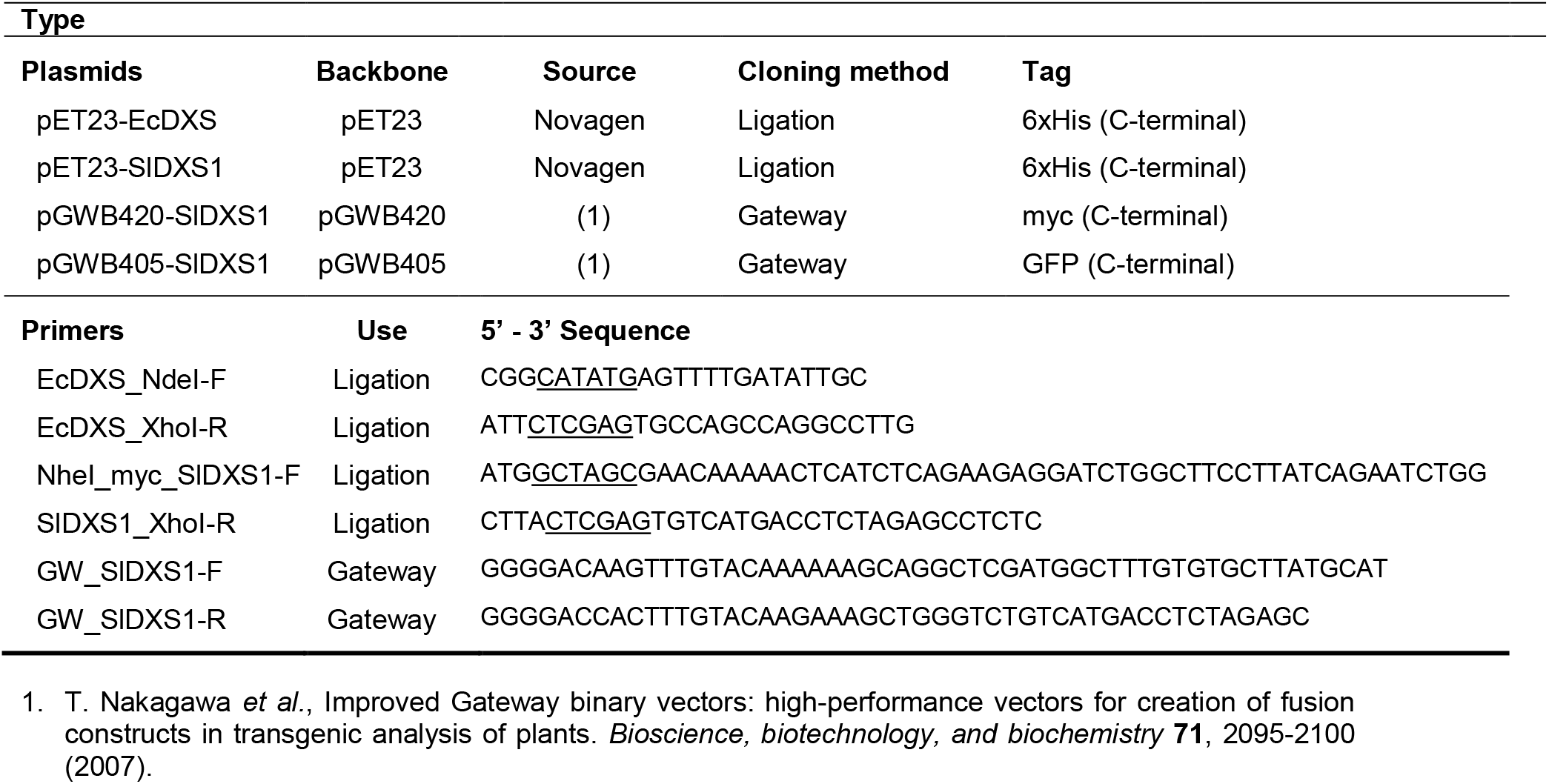
Plasmids and primers used in this study.

## Supplemental Figures

**Fig S1.**
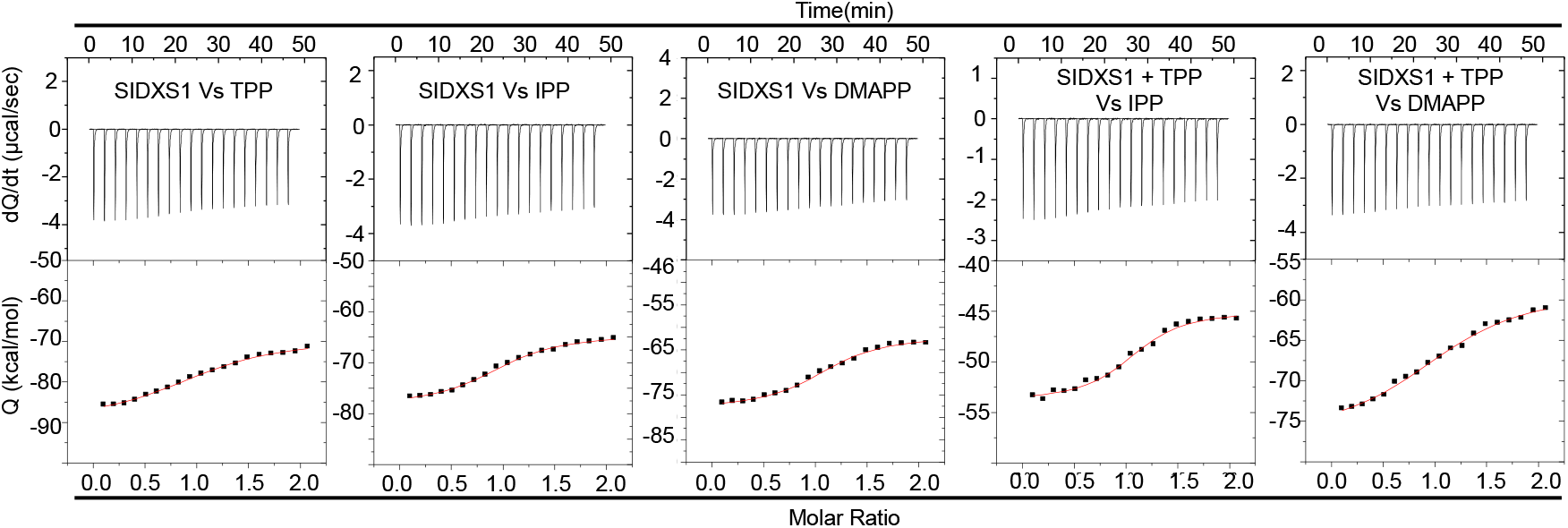
Analysis of SlDXS1 interaction with metabolite ligands. ITC plots were obtained from the titration of 20 μM SlDXS1 (either alone or with 100 μM TPP) with 200 μM TPP, IPP or DMAPP as indicated. Assays were performed at 25 °C in 50 mM HEPES, 150 mM NaCl buffer. The plots in the upper panel show the thermogram (raw thermal power as a function of time), and the plots in the lower panel show the binding isotherm (heat released per injection normalized per mole of ligand injected as a function of the molar ratio, [ligand]/[protein], in the calorimetric cell). The solid lines represent the best fits of the experimental data after non-linear lest-squares analysis using a single-site binding model.

